# Trends in scientific research on Environmental Education: A scientometric review

**DOI:** 10.1101/2020.09.14.297267

**Authors:** Leonardo Fernandes Gomes, Hasley Rodrigo Pereira

## Abstract

Given the need to understand the current approaches to Environmental Education (EE) in the world, we answer the following questions: (i) Have the studies directed to EE received attention from the scientific community? (ii) what are the trends in EE publications? In the past years, there has been an increase in the number of publications on EE. Brazil stood out in the number of publications, reflecting the concern of Brazilian researchers to promote sustainability and biodiversity maintenance. As for the approaches, the studies are broad, ranging from the influence of policies on environmental protection to the importance of reflection at a global level, proposing international agreements. However, regarding the EE application, given the multiple existing currents, it is worth emphasizing the importance of the teaching-learning process to take place critically so that there is no favoring of the contents promoted and supported by a hegemonic class.

## 1. Introduction

Given the increase in the world population, which increases the demand for space and resources to feed and shelter people (Crist et al., 2017), and the increasing human impacts resulting from human activities, several planetary boundaries have already been exceeded (Steffen et al., 2015). Due to the increase in urban populations and impacts on the environment, the need for Environmental Education (EE) emerges. The EE’s main objective is to teach people to be knowledgeable, aware and capable of proposing solutions to environmental problems (Stapp, 1969). Currently, the new perspective of EE consists of a paradigm shift that involves the construction of new public policies to build responsibility, diversity and solidarity among the agents involved. It is interpreted as an educational process that leads to knowledge of the environment from the perception of ethical values, political rules and social interaction that implies the management and use of nature (Guimarães, 2016; Sorrentino et al., 2005).

For some time, it was thought only interventional actions of EE would produce direct results on the reduction of environmental impacts. However, research has revealed that changes in attitudes of society are related to several factors that go beyond knowing the importance of environmental protection, as it involves a series of emotional, cognitive and cultural factors. Therefore, the need to associate EE with other knowledge areas, such as Scientific Education, where EE is looking for popular engagement on these issues, while other knowledge areas can bring proposals on how this engagement can bring environmental improvements with social participation (Wals et al., 2014).

EE can be a way to mitigate such effects by raising awareness among the population and decision-makers, who can intervene through public policies (Frantz & Mayer, 2014; Lorenzoni et al., 2007). However, this educational approach is challenging, since the relationship between man and nature is complex and involves cultural, religious, economic aspects, among others (Commoner & Egan, 2020; Jacobi, 2003). The change in society’s behavior concerning the environment requires, in addition to the transmission of knowledge, that environmental educators have an understanding of the culture, emotions and beliefs of the public they want to reach. Only in this way, it becomes possible to more effectively reach the population’s attitudes and behaviors (Pooley & O’Connor, 2000).

Also, human beings must recognize themselves as part of the world in which they live, so that they can contribute more effectively. In the Education of young people and children, the theme must be treated in a transversal way in the school subjects and go beyond the application in specific activities, in order to promote a more effective future progress in public policies (Corrêa & Barbosa, 2018). Thus, actions related to EE cover different forms of conception (chains), which can be strategically used according to the target audience and the educator’s conceptions (Sato & Carvalho, 2009).

### 1.2. Conceptions of Environmental Education

The expression “Environmental Education” became notorious in the second industrial revolution, mainly from the second half of the 20th century, when the use of natural resources increased (Bergstrom & Randall, 2016). Consequently, researchers and society have come to realize the need to sensitize the population of different places about ‘biophysical’ problems. During that same period, it became clear that man, culture and the environment are inseparable components. Therefore, actions of anthropic origin can have severe environmental impacts on contemporary society (Stapp, 1969). The movement gained strength from the Intergovernmental Conference on EE held in Tbilisi (USA), in 1977, which suggested strategic actions in favor of a new awareness about the value of natural resources, based on the construction of ideas in an interdisciplinary way, according to the complexity of this approach (Jacobi, 2003; Tblisi, 1977).

Despite this, over the years, multidisciplinary approaches to EE have emerged, and, in some cases, there has been a defense that certain understandings or propositions are more appropriate than others. With the breadth of these propositions, EE started to be subdivided into different currents (e.g., naturalistic, systemic, moral/ethics, holistic, eco-education, sustainability), according to different dominant aspects (e.g., sensory, cognitive, experimental, praxic, dialogistic, spiritual, affective and pragmatic) (Sato & Carvalho, 2009). Consequently, from these currents, different methodologies have emerged, in which we can highlight the most recent ones: holistic, bioregionalist, praxic, critical, feminist, ethnographic, eco-education and sustainability (Sato & Carvalho, 2009).

Thus, projects and actions aimed at effective EE can be highly diversified and make up different currents. Despite the specificities present in each approached chain, they are not mutually exclusive, which means they complement each other and can make environmental awareness more effective for all target groups (Sauvé, 2005a, 2005b).

### 1.4. Background and objectives

However, EE is often guided by the paths of globalization and public policies, which can be negative for the construction of knowledge on the subject, given that they are not starting from the problems raised by the academic community. These factors create the risk of EE being tied only to sustainable development, which leaves aside issues such as health, social justice and income distribution and ecosystem services in general (Costanza et al., 2017; Jickling & Wals, 2008).

Despite the importance of EE as a tool to raise citizens’ awareness of anthropic threats to the environment and the global discussions surrounding this topic, no work in the scientific literature points out the flow and trends of these debates. Thus, given the need to understand the current approaches to EE in the world, we seek to answer the following questions: (i) Have EE studies received attention from the scientific community? (ii) What are the trends in EE publications?

## 2. Methods

We carried out a scientometric review as an instrument for evaluating publications. Although there are numerous confusions among researchers about the differentiation between bibliometrics and scientometrics, it should be noted that scientometrics is much more associated with aspects that go beyond bibliometric indicators. Therefore, while bibliometrics is much more related to the number of publications, citations, areas of publication and authorship, scientometrics is more associated with aspects that permeate these indicators, such as public policies and other factors associated with literature production and the content of publications (Hood & Wilson, 2001).

### 2.1. Search for publications

The *Web of Science* database is one of the broadest concerning the coverage of scientific articles worldwide, in many cases being compared to Scopus, although there are different directions for publications indexed by each one (Martín-Martín et al., 2018; Mongeon & Paul-Hus, 2016). Therefore, to find publications related to Environmental Education, we performed an advanced search for titles and keywords in the main database of *Web of Science* with the following terms: (“environment* educ*”). We restricted the search years from 1991 (when publications abstracts started being indexed on the platform) to 2019. The use of quotation marks and asterisks allowed the search for expressions and the scope of derived terms, respectively. After this step, We extracted the publications’ data in text format (.*txt*).

We made the first evaluations through the HistCite ™ program, which allows the extraction of the number of publications per year, main authors, institutions and type of publication. These data were extracted in text format and made available in figures generated through the *R* program (R Core Team, 2016), package *ggplot2* (Wickham, 2016).

### 2.2. Network of words

To understand and to group the main approaches in publications on EE, we performed a mapping technique based on text files using the VOSviewer program (van Eck & Waltman, 2010). To perform the analysis and projection, those words that occurred in at least ten different publications (binary method) were selected. This projection technique uses the association force matrix. Closer words tend to occur simultaneously with greater frequency in publications. The larger dimensions of the circles refer to the total number of occurrences of the term. As a result, larger circles reflect words that occurred more frequently. *VOSviewer* also allows automatic clustering of words, so different colors refer to different groupings of words occurrences.

## 3. Results

We found a total of 3412 publications on Environmental Education. There was no significant growth in the number of publications until 2010 when the publications started to have an increasing number. As of 2015, there was a leap in the number of publications that went from 141, in 2014, to 348.

USA and Brazil stood out in the number of publications in comparison to the other countries (Figure 2.A). The top three institutions in the number of publications are Brazilian: Federal University of Rio Grande (FURG), University of São Paulo (USP) and Paulista State University (Unesp). Followed by the State University System of Florida, University of North Carolina, Cornell University, which are US institutions (Figure 2.B). Krasny ME was the author who published the most on the subject, followed by Kopnina H and Bogner FX (Figure 2.C).

**Fig. 1.**
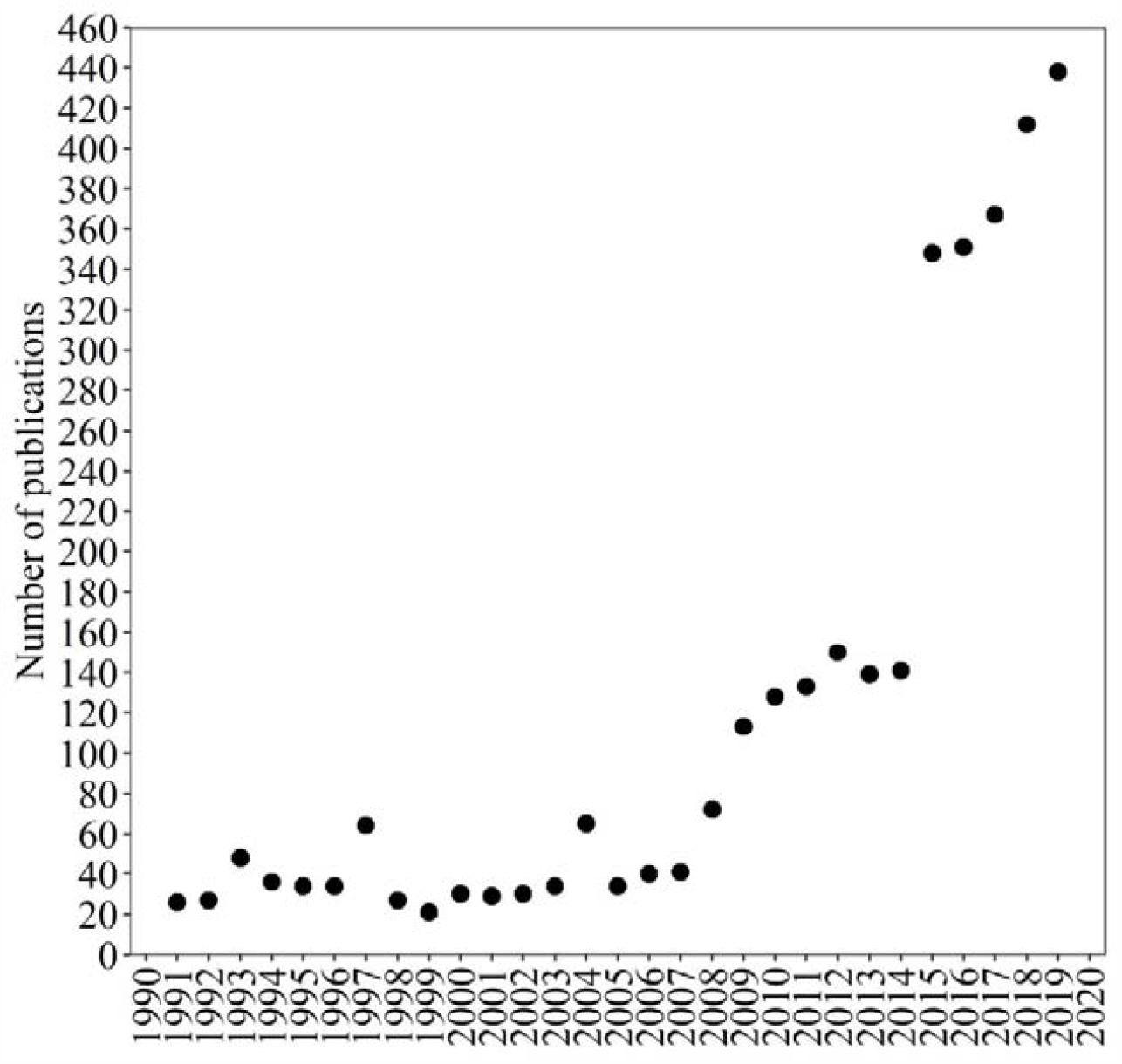
Number of publications per year on Environmental Education between 1991 and 2019

**Fig. 2.**
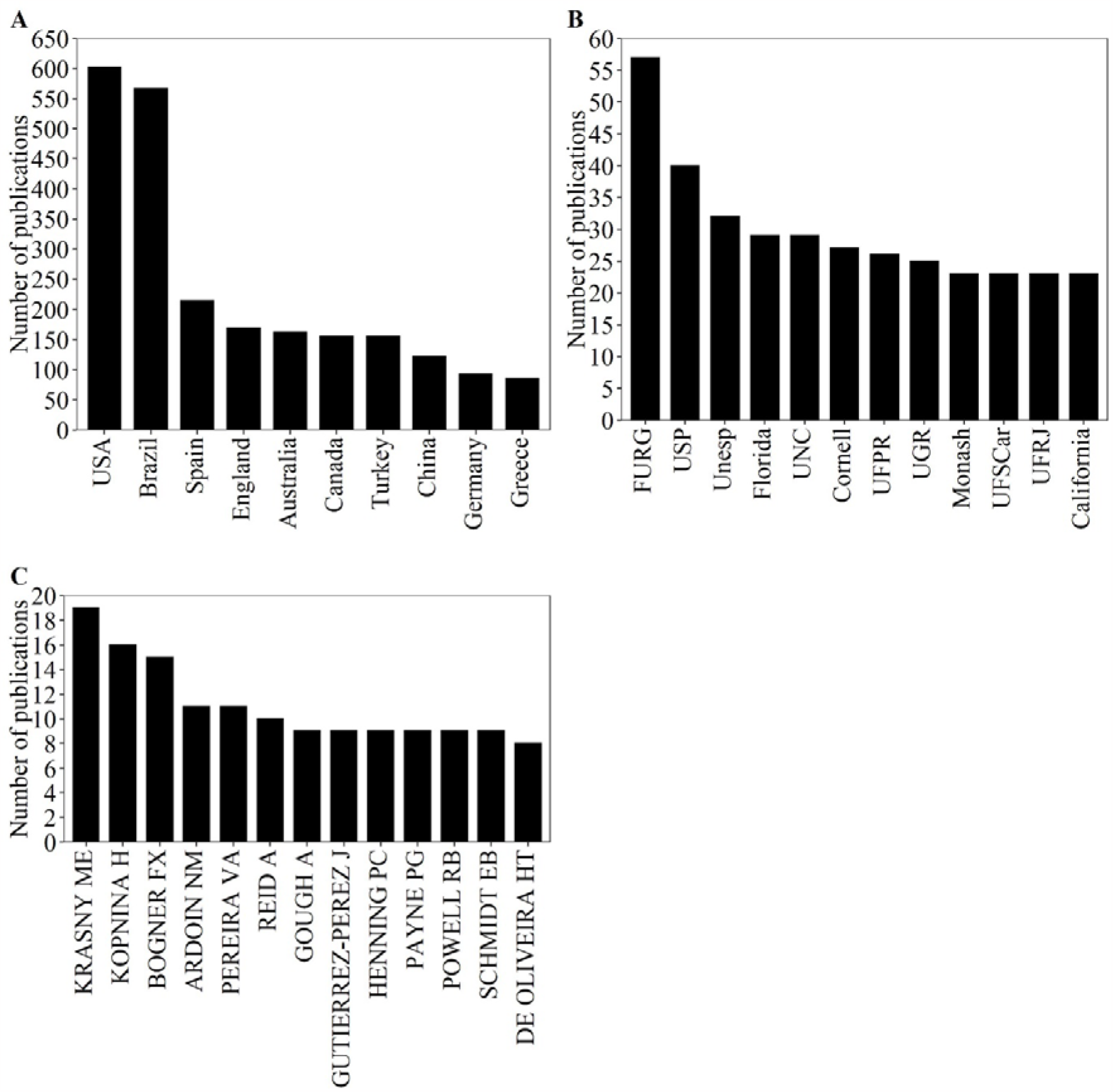
Number of publications on Environmental Education ranking by (A) country, (B) institution, and (C) author. Acronyms: USA = United States of America; FURG = Federal University of Rio Grande; USP = University of São Paulo; Unesp = Paulista State University; Florida = State University System of Florida; UNC = University of North Carolina; Cornell = Cornell University; UFPR = Federal University of Paraná; UGR = University of Granada; Monash = Monash University; UFSCar = Federal University of São Carlos; UFRJ = Federal University of Rio de Janeiro; California = University of California System

The main words used in the titles and abstracts of articles on EE formed six distinct groups. However, three groups were more prominent, with the following associated words (Figure 3): attitude, questionnaire, scale and environmental attitude (green); Possibility, Brazil, space, dialogue and discourse (blue); conservation, respondent, tourism, resident, visitor (red); College, higher education, pollution and industry (yellow).

**Fig. 3.**
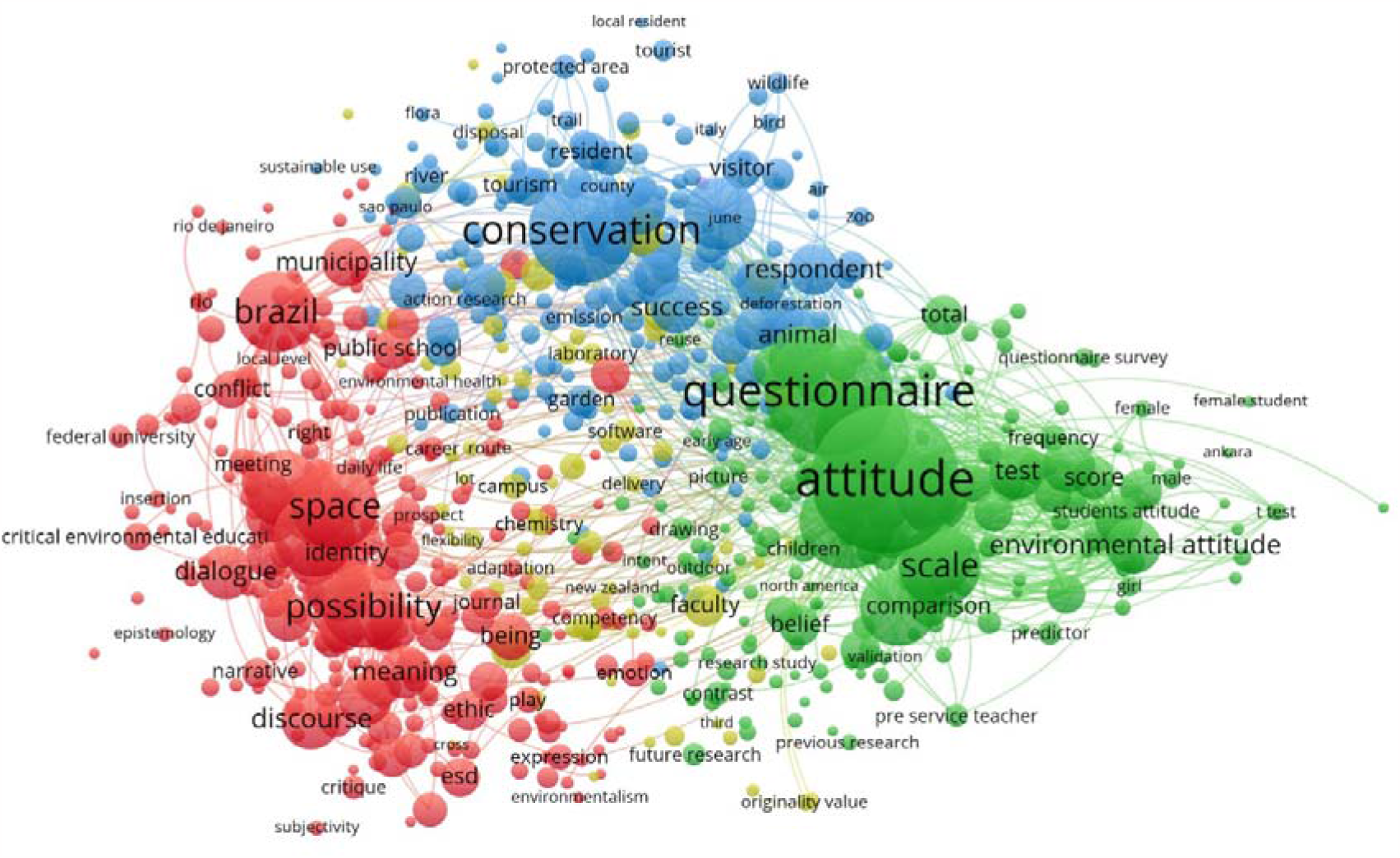
VOS projection of the main words used in publications on EE. The colors reflect different groupings of words. Words closest to each other tend to occur simultaneously, while the most distant tend not to occur simultaneously

## 4. Discussion

Seeking to understand publications on EE in the world, we found that there has been a high numerical growth over the past few years. That may be a result of increasing interest from the scientific community on the subject. The anthropic effects on natural ecosystems have been the subject of many controversies and socio-environmental conflicts. Therefore, there are several proposals for the resolution of such conflicts in national and international scenarios (Fisher & Sablan, 2018; Schultz et al., 2018). This scenario highlights the need for further research on ways to promote dialogues and alternatives to mitigate the effects of the anthropization of global ecosystems.

Not surprisingly, the US is first in the number of publications, considering that they have a large number of investments in education and research for different areas of knowledge (United Nations, 2015), which can influence the quantity and quality of publications. Brazil, as second in the number of publications, shows that researchers understand the dimension of the country as the shelter of several biomes in its territory and wide biological diversity, despite experiencing severe losses resulting from the expansion of anthropic activities and controversial decision-making policies regarding biodiversity protection (Bockmann et al., 2018; Ratter, 1997). Despite this, the resolution that established the national guidelines for EE in Brazil (BRASIL, 2012) may have been the factor responsible for the growing number of publications in that country and over the past few years.

The Federal University of Rio Grande (FURG), University of São Paulo (USP) and Paulista State University (Unesp) are three Brazilian universities among the leading institutions in the number of publications on EE. The number of publications shows that these universities are references on a global level and concentrate many researchers who deal with the subject. The universities State University System of Florida, University of North Carolina, Cornell University, institutions in the USA, also presented a high number of publications, which shows that most studies on the subject are developed at these universities.

In this sense, understanding the determining factors for certain institutions to publish more about EE can be a challenge. The interaction between social and environmental interfaces, aimed at promoting sustainability, involves a series of parameters, such as economy, EE and demographic aspects (Lehtonen, 2004). Therefore, the number of researches are related to the regional level of interests on the topic, the institutional motivation for promoting studies, the number of specialists in an institution and other factors, such as the number of graduate programs related to the topic (Kates, 2001).

Among the researchers who published the most on the subject, it is worth emphasizing Krasny, ME (Marianne E. Krasny). Krasny’s publications are mainly related to EE research in the face of the challenges posed by demographic growth and global environmental changes (Krasny, 2009). Some of Krasny’s prominent publications are also aimed at EE for young people (Krasny et al., 2015), where the value of social capital is evidenced in the integration between young people and adults in education and environmental awareness. Therefore, in this context, social capital refers to a set of social interactions that make public policies on EE effective and capable of modifying the lives of young people through various indicators, such as the reduction in the number of young pregnant women and reducing delinquency (Krasny et al., 2015).

Other studies that the same author participated in also highlighted that there are problems in defining the discourses, practices and results of the perspectives on EE, although all conceptions value human well-being and sustainable practices (Fraser et al., 2015). However, different perspectives can lead to a disagreement among researchers engaged in promoting advances in these studies, considering that some perspectives have a greater emphasis on the concern with the “non-human” nature, others on a greater affective connection of the human being with nature and others with public policies aimed at solving complex problems (Fraser et al., 2015). Therefore, it is important for the development of EE that all these perspectives are integrated. With this, there will be a reduction of conflicts and the emergence of different spaces that directly or indirectly promote sustainability and human well-being (Fraser et al., 2015).

Tied in the number of publications, the next authors in the number of publications raking were Kopnina, H (Helen Kopnina) and Bogner, FX (Franz X. Bogner). Bogner is linked to the University of Bayreuth, Germany. Kopnina is from the University of Cambridge (United Kingdom). Kopnina brought in one of her studies (Kopnina & Cocis, 2017) an approach directed to measures the environmental (ecocentric) attitudes of higher education students based on scales. The results were surprising, considering that the choice of courses directed to environmental areas did not directly reflect the attitudes of the students evaluated. Also, the same study mentions that despite the great concern with ecocentric attitudes, defended by EE, there is also a need to assess how sustainable development objectives can reduce problems related to income distribution. Therefore, EE must cross the attitudes of individuals and their actions towards life in society with a sustainable bias. However, it must reach the means of production so that they are consistent with the reduction of poverty, improvement in the income distribution and the mitigation of competition for resources effects (Kopnina & Cocis, 2017).

Bogner’s studies are mainly related to the results of implementing practical activities on improving cognitive knowledge (Dieser & Bogner, 2016) and how obtaining knowledge related to nature interferes with a more ecological way of life (Roczen et al., 2014). Therefore, through activities outside the classroom, students have the opportunity to put into practice what was learned. Also, they can overcome prejudices concerning contact with animals (Dieser & Bogner, 2016). Therefore, these activities have great potential for transforming students’ awareness of the environment and the search for attitudes more consistent with sustainable development.

### 4.1. Main approaches

Four main approaches were outlined in publications on EE. The first, strongly related to environmental protection. EE can promote ways to achieve environmental protection effectively, Mainly through interaction between people and nature, which promotes protective behavior in individuals (Frantz & Mayer, 2014). Very close to the first approach, another highlight was the attitudes related to EE, which promoted the development of assessments of the direct and indirect effects of education, such as the influence of EE on the parents of children who learned about the topic at school and disseminated attitudes such as recycling materials at home (Evans et al., 1996). There is also debate about the importance of emotions and beliefs that, in many cases, can be more valuable for changing attitudes towards the environment than in-depth knowledge on the topic (Pooley & O’Connor, 2000). On the other hand, some mistaken traditional beliefs and knowledge about environmental issues can be restructured based on practical actions by environmental teachers/educators (Hofstatter et al., 2016).

The groups of approaches that deal with reflection, proposals, ideas, the world and agendas were also strongly related to each other. Faced with the emergence of environmental problems (Crist et al., 2017), the world has sought different ways to improve this scenario. Among them, the establishment of international agreements aimed at the common good, as is the case of the Paris Agreement, intending to mitigate the effects of global warming and the millennium goals in favor of sustainable development (Griggs et al., 2013; United Nations, 2015).

### 4.2. Pedagogical trends and Environmental Education currents

Education is linked to social and political contexts and, therefore, follows trends that differ regarding the role of the school, teaching content, methods and the relationship between teachers and students (Libaneo, 1983). In the same way, EE also follows trends that are distributed in 15 different currents (Sato & Carvalho, 2009) that differ concerning the prevailing conceptions of the environment, objectives and approaches. In this context, it is important to emphasize that no current is superior or should receive more attention from environmental educators.

However, currently, the critical-social trend of the content has been highlighted in countries such as Brazil, one of the main players in research on EE. In this trend, the school role and the content must be linked to the social realities of the students with whom we are interacting, in addition to providing a more critical analysis concerning the contents that are linked to the teaching-learning process, so as not to favor a hegemonic culture (Libaneo, 1983). Therefore, the current context requires that, regardless of the current to which the teaching-learning process of EE is linked, there is a critical stance to what is being transmitted, where the Teacher is the mediator of the teaching-learning process and promotes a stance among students that does not favor the development of a model that favors hegemony, where historically the dominant classes uncritically determine what should be learned and taught (Libaneo, 1983).

## 5. Conclusions

Over the past few years, there has been an increasing number of publications on Environmental Education, which shows that the scientific community has been interested in promoting attitudes, debates and evaluations related to the theme in different spheres of society. Also, the USA and Brazil stood out in the number of publications on the topic, which reflected their researchers’ concern with promoting sustainability and maintaining biodiversity.

As for the approaches, it is important to emphasize that the studies on EE have their foundation in different areas of knowledge and, therefore, they can address topics ranging from the influence of EE policies on environmental protection to others that deal with the importance of reflection at the global level, with the proposition of international agreements. From that, we suggest that the global scientific community pays more and more attention to EE, so that there are constant advances in favor of the formation of sensitive citizens regarding the problematic degradation of the environment and, thus, these can contribute to the conservation of natural resources and ensure them for the next generations.

It should also be noted that despite the focus of some studies on EE in a broader context, knowledge and awareness can start at more local scales. In this case, it is important to emphasize more regionalized public policies and classroom teaching since the early years. From the integration of these local and global public policies, there is a greater chance of success towards more sustainable social practices. However, it is worth emphasizing the importance that the teaching-learning process takes place in a critical way so that there is no favoring of the content promoted and supported by a hegemonic class.

## Disclosure statement

The author(s) declared no potential conflicts of interest with respect to the research, authorship, and/or publication of this article

## Notes

### Competing Interest Statement

The authors have declared no competing interest.

### Summary of Updates

This version was adjusted to the journal: "Educacao em Contexto"

